# Familial Mutations and Post-translational Modifications of UCH-L1 in Parkinson’s Disease and Neurodegenerative Disorders

**DOI:** 10.1101/094953

**Authors:** Yun-Tzai Cloud Lee, Shang-Te Danny Hsu

## Abstract

Parkinson’s disease (PD) is one of the most common progressive neurodegenerative disorders in modern society. The disease involves many genetic risk factors as well as a sporadic pathogenesis that is age- and environment-dependent. Of particular interest is the formation of intra-neural fibrillar aggregates, namely Lewy bodies (LBs), the histological hallmark of PD, which results from aberrant protein homeostasis or misfolding that results in neurotoxicity. A better understanding of the molecular mechanism and composition of these cellular inclusions will help shed light on the progression of misfolding-associated neurodegenerative disorders. Ubiquitin carbonyl-terminal hydrolase L1 (UCH-L1) is found to co-aggregate with α-synuclein (αS), the major component of LBs. Several familial mutations of UCH-L1, namely p.Ile93Met (p.I93M), p.Glu7Ala (p.E7A), and p.Ser18Tyr (p.S18Y), are associated with PD and other neurodegenerative disorders. Here, we review recent progress and recapitulate the impact of PD-associated mutations of UCH-L1 in the context of their biological functions gleaned from biochemical and biophysical studies. Finally, we summarize the effect of these genetic mutations and post-translational modifications on the association of UCH-L1 and PD in terms of loss of cellular functions or gain of cellular toxicity.

## 1. INTRODUCTION

### 1.1 ETIOLOGICAL FACTORS AND PATHOLOGICAL PROTEINS IN PD

More than 90% of PD cases are sporadic in origin and progress with age over 65 years old. Two pathological hallmarks in the diagnosis are selective degeneration of dopaminergic neurons in substantia nigra (SN; located in the midbrain) and the presence of an intracellular protein inclusion, Lewy bodies (LBs). Although the etiological factors of sporadic PD remain elusive, several PD-monogenetic forms and susceptible genetic risk factors have been identified, including parkin, PTEN-induced protein kinase 1 (PINK1), DJ-1, lysosomal P-type ATPase (ATP13A2), leucine-rich repeat kinase 2 (LRRK2) and presynaptic nerve terminal protein SNCA (α-synuclein precursor) (Table 1) [1, 2]. Most of these PD risk factors are involved in regulation of a variety of physiological roles that contribute to nigral dopaminergic cell death *via* mitochondrial dysfunction, abnormal response to oxidative stress and inflammation, and altered protein homeostasis [3]. In addition, a number of studies have reported associated non-genetic environmental factors or endogenous toxins such as 1-methyl-4-phenyl-1, 2, 3, 6-tetrahydropyridine (MPTP) neurotoxin, rotenone, paraquat (1, 19-dimethyl-4, 49-bipyridinium dichloride), and 6-hydroxydopamine (6-OHDA), which can induce PD-like symptoms in mice [4-7]. All these molecules are oxidative stress inducers that directly or indirectly modify biomolecules, thereby disrupting their normal cellular functions.

**Table 1.** Summary of genetic PD risk factors and their biological functions.

| Functional roles | Protein Name | Gene Loci | Biological functions | Familial mutations | Reference |
| --- | --- | --- | --- | --- | --- |
| **Mitochondria integrity maintenance** | Parkin | PARK2 | RING-between-RING type Ub E3 ligase | About 170 mutations | [97] |
|  | PINK1 (PTEN induced putative kinase 1) | PARK6 | Serine/threonine protein kinase | About 70 mutations | [98] |
|  | LRRK2 (Leucine-rich repeat kinase 2/Dardarin) | PARK8 | GTPase/Protein-Protein interaction/protein kinase | p.G2019S, p.G2385R and p.R1628P | [99] |
| **Proteostasis maintenance** | UCH-L1 (Ubiquitin C-terminal hydrolase L1) | PARK5/PGP9.5 | Deubiquitinase/mono-Ub stabilizer/Ub ligase | p.S18Y, p.I93M and p.E7A | [8] [9] |
| **Oxidative stress response** | DJ-1 | PARK7 | Antioxidative stress reaction/chaperone/mitochondrial regulation | About 23 mutations | [100] |
| **Presynaptic nerve terminal protein** | SNCA (α-synuclein precursor) | PARK1/PARK4 | Trafficking neurotransmitters | p.A30P, p.E46K and p.A53T | [101] |
| **Lysosomal functions** | ATP13A2 | PARK9 | Lysosomal P-type ATPase | p.F182L, p.G504R | [102] |
| | GBA | Not assigned | β-Glucocerebrosidase | p.L444P, p.D406H | [103, 104] |

Ubiquitin carbonyl-terminal hydrolase L1 (UCH-L1), which is also known as protein gene product 9.5 (PGP9.5), has been suggested to be a risk factor of PD, and indeed, a number of familial mutations have been reported to be associated with increased risk of PD [8]. In addition to the familial mutations in UCH-L1, post-translational modifications (PTMs) of UCH-L1 may play an important role in PD and other neurodegenerative diseases [9, 10]. Nevertheless, despite its abundance in neuronal cells, the molecular basis of the functional implications of the familial mutations and PTMs of UCH-L1 in the context PD has not been firmly established [11]. In this review, we shall discuss the current structural, biochemical and functional understandings of the association of UCH-L1 with PD and neurodegenerative diseases.

### 1.2 BIOLOGICAL FUNCTIONS OF UCH-L1

UCH-L1 is one of the most abundant proteins, expressed specifically and highly in brain neurons, accounting for about 2% of total soluble protein content [12-14]. It is a member of the human ubiquitin carbonyl-terminal hydrolases (UCHs), one of the four families of deubiquitinases (DUBs), namely UCHs, ubiquitin-specific proteases (USPs), Machado-Josephin domain proteases (MJDs) and ovarian tumor proteases (OTUs). Together, there are in total 95 DUBs in the human genome [15]. Many DUBs are implicated in malignant transformation because of their important role in regulating cell proliferation [16]. Indeed, UCH-L1 has been suggested to be an oncogene for various forms of cancer [17], potentially because of its involvement in the ubiquitin-proteasome system (UPS) [18]. For example, UCH-L1 was found up-regulated in colorectal cancer [19] and breast cancer [20], and has been suggested to promote the development of lymphoma [21, 22] and prostate cancer [23]. UCH-L1 is also implicated in the development of lung carcinoma [24]. Indeed, suppression of the DUB activity of UCH-L1 by small molecule inhibitor promotes proliferation of H1299 lung cancer cell line [25]. Using the same inhibitor, a recent study using cell cultures has demonstrated that UCH-L1 promotes metastases as a deubiquitinating enzyme for HIF-1α [26]. The underlying molecular mechanism of UCH-L1-associated metastasis may be linked to its association with adhesion complexes that promote cell migration and anchorage-independent growth [27]. Recently, UCH-L1 was found to regulate the activities of a number of cyclin-dependent kinases (CDKs), which may in turn enhance cancer cell proliferation independent of its DUB activity [28]. Conversely, in prostate cancer and HeLa cell, UCH-L1 is a potential tumor suppressor whose promoter is usually silenced by methylation in these cancer cells, but not in normal cells [29, 30].

In the context of neurobiology, intragenic deletion of UCH-L1 results in the gracile axonal dystrophy (*gad*) mouse that shows early-onset sensory ataxia followed by motor ataxia [31]. The UCH-L1 null mutation in mice also leads to progressive paralysis and premature death [32]. UCH-L1 and its homolog UCH-L3 appear to have redundant functions [33]. Double knock-out of UCH-L1 and -L3 in mice led to weigh loss and early lethality due to dysphagia, lending support to the hypothesis that UCH-L1 and -L3 have both separate and overlapping functions in the maintenance of neurons [33]. Overexpression of α-synuclein (αS) in *gad* mice results in early-onset motor deficits and forebrain astrogliosis [34], although it dose not support a clear role in regulating αS-induced toxicity. At the molecular level, it has been shown that down-regulation of UCH-L1 in mouse neuron and embryonic cell (MES) can decrease the level of monoubiquitin (monoUb) without significant alteration of the mRNA levels of Ub precursor genes. Hence UCH-L1 has been proposed to be a monoUb stabilizer [35]. Despite being a relatively inefficient DUB, the DUB activity as well as the monoUb stabilization capacity of UCH-L1 are implicated in regulating synaptic functions at glutamatergic synapses [32, 36]. Specifically, treatment of N-methyl-D-aspartate (NMDA), which is a neuron transmitter, resulted in increased activation of UCH-L1 and increased levels of monoUb in cultured hippocampal neurons. In contrast, pharmacological inhibition of UCH-L1 activity using UCH-L1-specific inhibitor LDN-5744 resulted in a decreased level of monoUb and slowed down proteasome-mediated degradation [36]. Interference of Ub pool in neurons was accompanied by defects in synapse structures, including decreased spine density, increased spine size, and increased accumulation of pre- and post-synaptic proteins [36]. Taken all together, these results suggest that homeostasis of monoUb level and activation of UCH-L1 in neurons is critical for maintaining synaptic functions and that loss of the DUB activity of UCH-L1 has a direct impact on synaptic integrity and plasticity.

As far as a DUB is concerned, UCH-L1 is not an effective DUB compared to other UCHs such as UCH-L3, which is ubiquitously expressed in human tissues [37]. It cannot hydrolyse isopeptidic linkages within polyubiquitin (poly-Ub) chains *in vitro* because of the short active-site crossover loop that prevents access of large ubiquitinated substrates into its catalytic site; by grafting the crossover loop of UCH-L5 (also known as UCH37), which is six-residues longer than that of UCH-L1, the chimeric UCH-L1 could then bind to and hydrolyse Lys48-linked di-ubiquitin (diUb) [38]. Although UCH-L1 cannot hydrolyse poly-Ub chains, it has been reported to be involved in the process of preubiquitin (preUb) with a C-terminal extension in order to complete maturation of monoUb and of small ribosomal proteins (Figure 1) [39]. The monoUb stabilizing function of UCH-L1 may be associated with its strong binding affinity for Ub, which has a dissociation constant *K_d_* of tens of nanomolarity (nM) [40]. UCH-L1 covers nearly half of the solvent accessible surface area of ubiquitin (Ub) upon complex formation, and this may therefore protect Ub from proteolysis as a way to stabilize monoUb.

**Figure 1.**
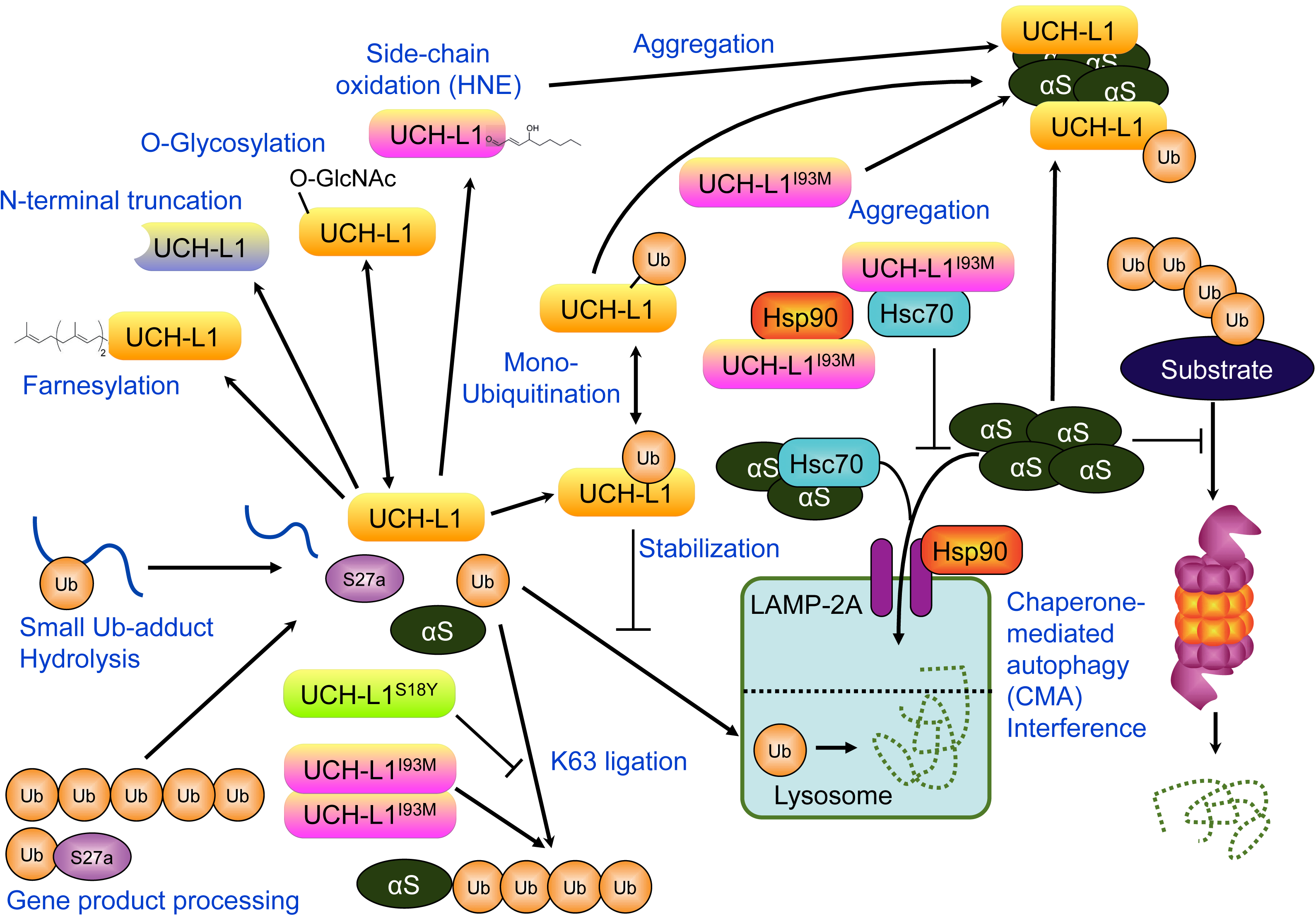
Schematic overview of UCH-L1 functions and regulations. The functions of UCH-L1 were suggested to involve in hydrolysis of ubiquitin C-terminal adducts and maturation of monoUb. The clearance of α-synuclein (αS) and other misfolding proteins can be achieved by the UPS and/or CMA. UCH-L1^I93M^ exhibits increased aggregation propensity compared to other UCH-L1 variants while all UCH-L1 variants become aggregation-prone as a result of down-regulation of UPS. UCH-L1^I93M^ exhibits aberrant interactions with HSP90 and HSC70 that are involved in CMA. Interference of CMA can therefore lead to accumulation of misfolded proteins.

Being a DUB and a potential monoUb stabilizer, UCH-L1 plays an important role in the UPS. Ardley *et al*. used immunofluorescence microscopy to observe a higher amount of PD-associated I93M variant (UCH-L1^I93M^) aggregates in COS-7 cell compared to wild-type and the p.S18Y variant even without perturbing the UPS. Introduction of a proteasome inhibitor (MG132) lead to overall increase of all UCH-L1 variants and co-localized punta of UCH-L1, parkin and αS [41]. There results showed that UCH-L1 is implicated in UPS and that down-regulation of UPS will results in aggregation of several PD risk factors, including UCH-L1. Furthermore, the naturally-occurring autosomal-dominant p.I93M variant, UCH-L1^I93M^, exhibits increased aggregation propensity without UPS interference. While UPS is the major pathway for clearance of misfolded proteins for proteostasis, clearance of αS aggregates can also be achieved by chaperone-mediated autophagy (CMA), which is independent of UPS [42]. It has been demonstrated that UCH-L1^I93M^ exhibits aberrant molecular interactions with heat shock proteins Hsp90 and Hsc70, which are involved in CMA, thereby reducing αS clearance [42]. Such aberrant interactions are shared by post-translationally modified UCH-L1 due to oxidative modifications by a reactive lipid metabolite, 4-hydroxyl-(2E)-nonenal (HNE), which could be implicated in sporadic PD [43]. PTMs of UCH-L1 are prevalent, and these include O-glycosylation [44], farnesylation [45], monoubiquitination [46], nitroxylation [47] and the N-terminal truncated isoform (Figure 1) [48]. The biological implications of these PTMs will be discussed in the following sections.

The increased aggregation propensity of UCH-L1 variants, which may lead to the formation of toxic species, may be more biomedically relevant. DUB activity-independent functions of UCH-L1 may also play an important role regulating physiological functions. Recently, Wada and co-workers showed that UCH-L1 could regulate cell proliferation *via* physical interactions with tubulins and CDKs that are involved in the cell cycle [28]. UCH-L1 p.R63A and p.H185A mutants showed increased interactions with α- and β-tubulin. Importantly, all these mutants showed stronger binding to LAMP-2A, a lysosome receptor pivotal in CMA (Figure 1) [42]. Of note, Arg63 and His185 are highly conserved in the UCH family (Figure 2).

**Figure 2.**
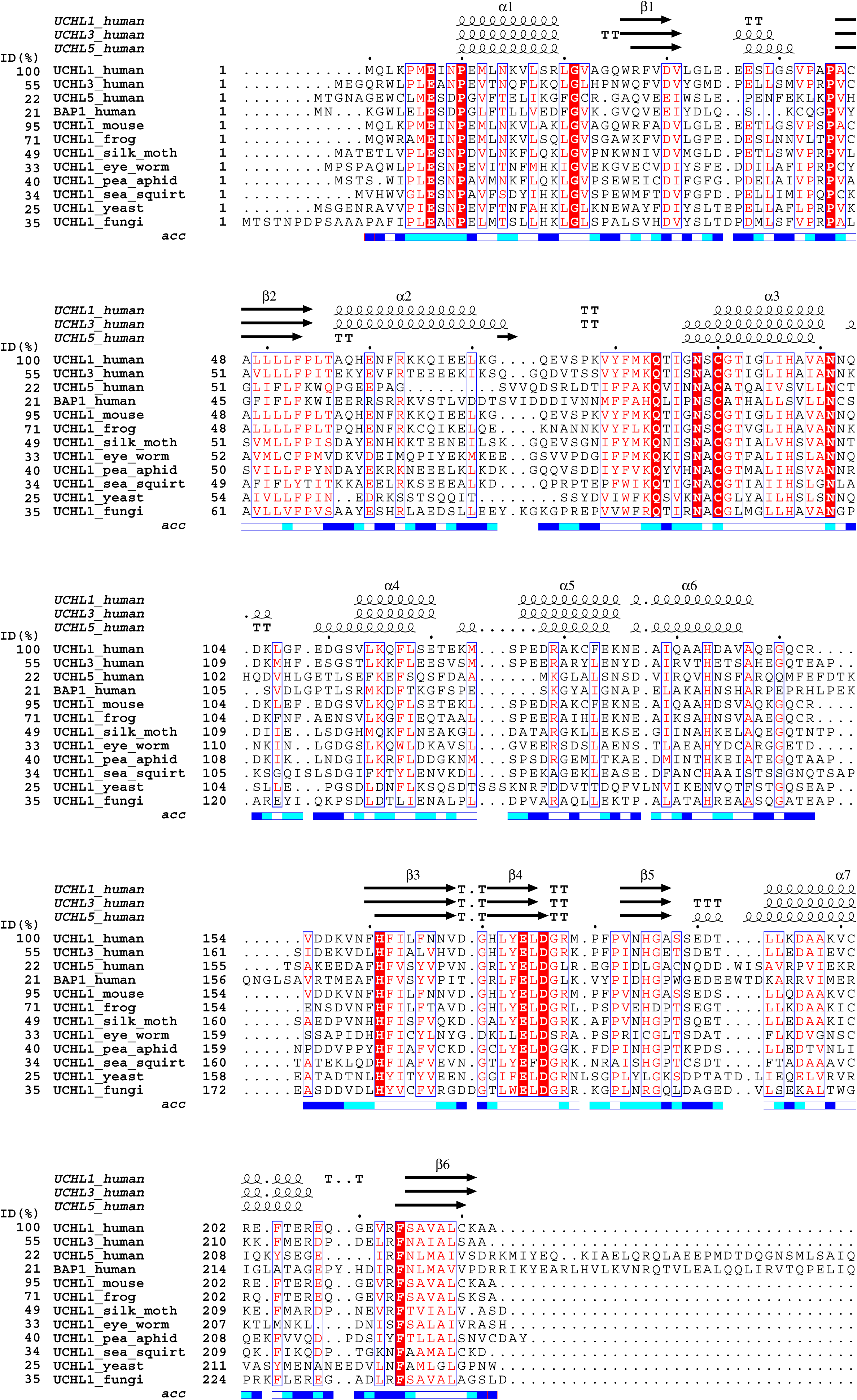
Sequence alignment of the catalytic domains of selected UCHs. The sequences were selected from the BLAST Assembled RefSeq Genomes. The sequence alignment and coloring were prepared by ESPript 3.0. The gene IDs are shown for each entry followed by the names of the proteins and species. The solvent accessibility (acc) of individual residues is colored black, grey or white at the bottom of the sequence alignment based on the crystal structure of UCH-L1 (PDB ID: 2ETL). Residues that are located in β-strand, α-helix or turn in the crystal structures of UCH-L1, -L3 and -L5 are shown in arrows, helices or T, respectively, in the top panels.

Unlike most DUBs, UCH-L1 has been reported to exhibit an unusual Ub ligase activity that is associated with the formation of non-covalent dimers and poly-ubiquitination of αS [49]. As well, the p.S18Y variant of UCH-L1 (UCH-L1^S18Y^) reduces the ability for dimerization, which in turn suppresses the ligase activity of UCH-L1^I93M^ *in trans*. Note that the Ub ligase activity was only observed when Ub-AMC was used together with a Lys-to-Arg Ub variant to form a linearized Lys63-linked diUb (Figure 1). The molecular machineries of UCH-L1 dimerization as an ubiquitinyl ligase in regulating biological function remain elusive to date. If dimeric UCH-L1 transfers acyl monoUb from the catalytic Cys90 to the side-chain of Lys63 of the proximal Ub, an alternative proximal Ub binding site may be introduced upon UCH-L1 dimerization. The molecular basis of such a process remains to be established. Recently, a study showed that dimeric UCH-L1 overexpressed in lymphoid and epithelial cells could be coimmunoprecipitated with α- and β-tubulin, providing biochemical evidence of the involvement of dimeric UCH-L1 in regulating microtubule dynamics *in vivo* [50]. Additionally, dimerization of UCH-L1 was also observed *in vitro* by small-angle neutron scattering and analytical ultracentrifugation [49, 51].

### 1.3. BIOCHEMICAL AND STRUCTURAL PROPERTIES OF UCH-L1

UCH-L1 is a single domain protein with 223 amino acids in length. It belongs to a α/β fold with five β-strands inside the hydrophobic core, which is surrounded by seven α-helices (Figure 3). The tertiary structure and catalytic residues of the UCH domains are highly conserved (Figures 2 and 3b) [52]. While most DUBs contain a Ub interacting motif (UIM) in addition to their catalytic domains to achieve substrate specificity and selectivity [53], UCHs utilize the same domain for Ub binding and hydrolysis. Indeed, structural mapping of the sequence conservation of the UCH family shows that the most highly conserved residues are located at the Ub binding interface (Figure 2 **and** 3a), which supports the functional importance of monoUb stabilization upon UCH binding. As mentioned in the previous section, UCH-L1 contains a flexible crossover loop, corresponding to residues 150 to 160, which is the shortest among the four human UCHs (Figure 2). The length and sequence composition of the crossover loop may affect the internal dynamics and more importantly substrate specificity.

**Figure 3.**
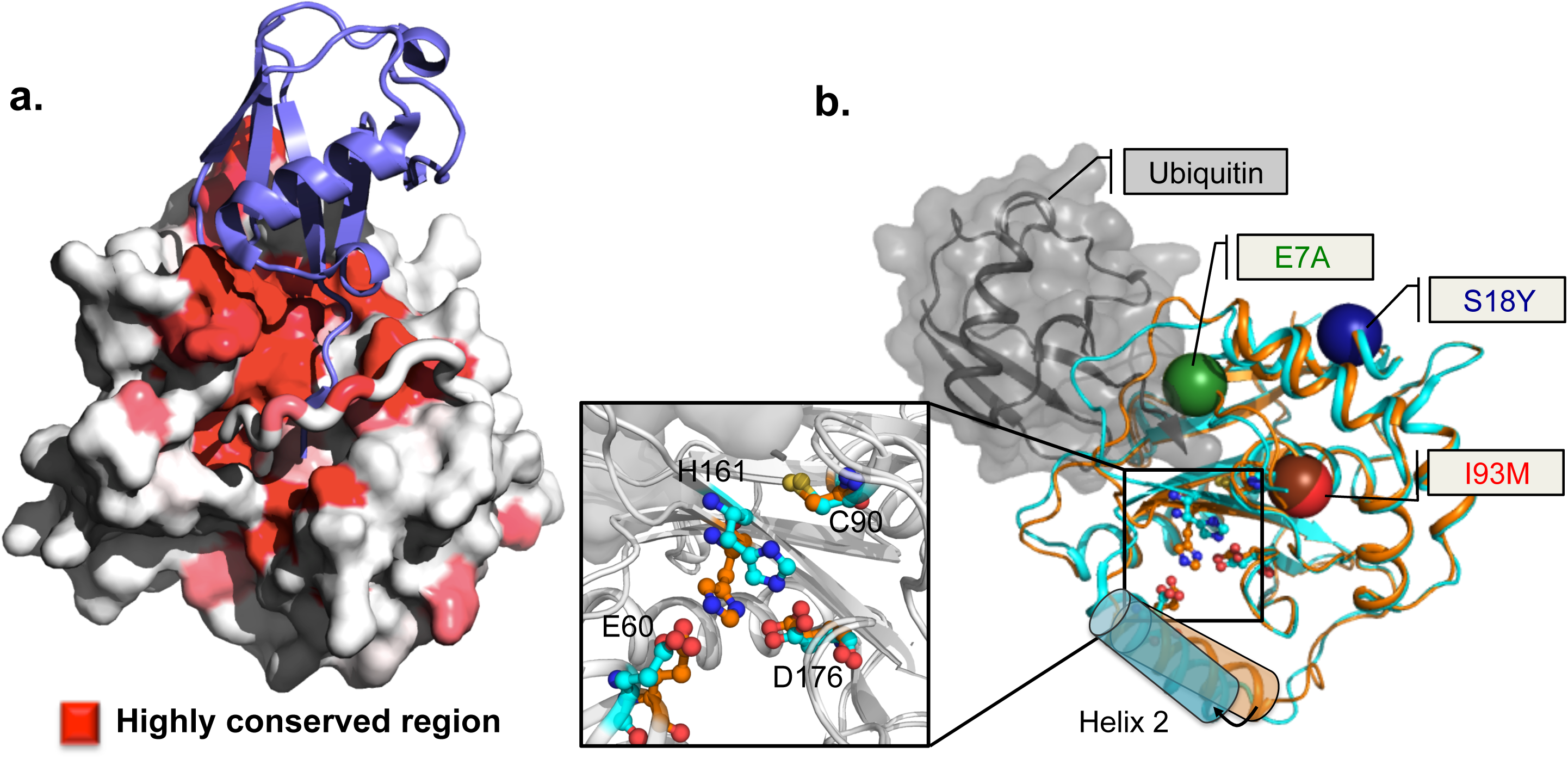
Structural mapping of sequence conservation and sites of neurodegeneration-associated mutations of UCH-L1. **a.** Structural mapping of the highly conserved residues onto the structure of ubiquitin-bound UCH-L1, shown in surface representation. The residues are color-ramped from dark grey to white according to their sequence conservation from high to low, respectively. The crossover loop of UCH-L1 is shown in tube representation. Ubiquitin is shown cartoon representation and is colored in grey. **b.** Structural mapping of the disease-associated mutations. The mutations sites of p.E7A, p.S18Y and p.I93M are shown in spheres and their identities are labeled respectively. The crystal structure of apo UCH-L1 (dark grey) is superimposed with that of ubiquitin-bound UCH-L1 (light grey) (PDB entries: 2ETL and 3KW5, respectively). While most of the structures of apo- and ubiquitin-bound UCH-L1 are very similar, ubiquitin binding does induce significant conformational change in the relative orientation of the helix 2 of UCH-L1 (indicated in rectangular boxes with an arrow showing the direction of the movement induced by ubiquitin binding). Ubiquitin is shown in semi-transparent grey surface representation with cartoon representation within. Inset: Enlarged view of the catalytic triad. The side-chains of Cys90 (C90), His161 (H161), and Asp176 (D176) are shown in ball-and-stick representation. Glu60 (E60) is a conserved residue that is involved in hydrogen bonding with H161. The side-chains of the apo- and ubiquitin-bound forms are shown in dark and light grey, respectively.

To characterize the enzyme mechanism of UCHs, Ub carbonyl-terminal 7-amido-4-methylcoumarin (Ub-AMC) has been developed as a fluorogenic substrate for UCHs [54]. The use of Ub-AMC has enabled detailed characterization of the enzyme mechanisms of UCH-L1, -L3 and -L5 [37, 55,56]. UCH-L1 turns out to be an inefficient DUB despite its abundance in brain neurons (Table 2). This therefore raised the question of whether the loss of the DUB function of UCH-L1 could potentially be compensated by other DUBs, such as UCH-L3, and hence the loss of function may not be responsible for the disease associations of the UCH-L1 variants. Regardless of the issue of functional redundancy of DUBs, the effects of pathogenic mutations as well as the functional role of conserved residues around the catalytic sites of UCHs have been elucidated with the aid of Ub-AMC (Table 2). Furthermore, a number of potent UCH inhibitors have been developed with the aid of a high-throughput screen based on similar used of Ub-AMC [25, 57, 58]. Of special note here, the use of Ub-AMC should be treated with caution, as it is not a real physiological substrate for UCHs and other DUBs. To address this issue, isopeptide linkage-conjugated ubiquitinated peptides have been developed as models for physiological substrates of DUBs [59, 60].

**Table 2.** **Enzyme kinetics parameters of UCH-L1 variants using Ub-AMC as the model substrate.** *k*_cat_ is the enzymatic turnover number of UCH-L1 derived from *V*_max_ of Michaelis-Menten steady-state enzyme kinetic model. *K*_M_ is Michaelis-Menten constant that is associated with substrate-binding specificity with UCH-L1.

| UCH variants | $k_{cat}$ ($s^{-1}$) | $K_M$ ($\mu M$) | $k_{cat}/K_M$ $10^6$ ($M^{-1} s^{-1}$) | Reference |
| --- | --- | --- | --- | --- |
| UCH-L1<sup>WT</sup> | 0.010 | 0.034 | 0.31 | [37] |
| UCH-L1<sup>WT</sup> | 0.02 | 0.040 | 0.50 | [105] |
| UCH-L1<sup>WT</sup> | 0.035 | 0.047 | 0.74 | [55] |
| UCH-L1<sup>WT</sup> | 0.174 | 0.12 | 1.45 | [43] |
| UCH-L1<sup>Q84A</sup> | 0.0011 | 0.056 | 0.02 | [55] |
| UCH-L1<sup>I93M</sup> | 0.079 | 0.11 | 0.72 | [43] |
| UCH-L1<sup>S18Y</sup> | 0.196 | 0.136 | 1.44 | [43] |
| UCH-L1<sup>E7A</sup> | 0.017 | - | < 0.01 | [9] |
| UCH-L3 | 5.9 | 0.02 | 277.3 | [105] |
| UCH-L3 | 18.6 | 0.077 | 241.4 | [55] |
| UCH-L5<sup>1-240</sup> | 33.67 | 21.5 | 15.7 | [55] |

**Table 3.** Effects of PTMs on UCH-L1

| Type of modification | Modification site | Effects of modification | Reference |
| --- | --- | --- | --- |
| Farnesylation | Cys220 | Membrane association and promotion of αS accumulation | [45] |
| Monoubiquitination | Lys4 and Lys157 | Reduced ubiquitin binding | [46] |
| N-terminal truncation | Eleven residue-truncation at the N-terminus | Prevent oxidative damage by Cys90-linked dimer associated with mitochondria. | [48] |
| 4-hydroxyl-(2E)-nonenal (HNE) | Cys90 and Cys152 | Protein aggregation and unfolding | [43] |
| 1-(3',4'-dihydroxybenzy)-1,2,3,4-tetrahydroisoquinoline (3',4'DHBnTIQ) | Cys152 | Protein aggregation and unfolding | [94] |
| Cyclopentenone prostaglandin | Cys152 | Protein aggregation and unfolding | [93] |

### 1.4. GENETIC ASSOCIATIONS BETWEEN UCH-L1 AND PD

Despite its lower DUB activity, UCH-L1 is closely associated with neurodegenerative diseases [61]. The expression level of UCH-L1 was found to be elevated in the hippocampal proteome in Alzheimer’s disease (AD) [62]. UCH-L1 is known as *PARK5*, which reflects its genetic association with PD [63-65]. Immunohistochemistry staining showed that UCH-L1 frequently co-aggregates with αS in LBs [66]. The *gad* mouse has an intragenic deletion of the UCH-L1 gene, which leads to typical neurodegenerative phenotypes, including sensory and motor ataxia in addition to the “dying-back” type of axonal degeneration [67] and formation of protein inclusions in nerve terminals [31]. In cell cultures, the aggregation of UCH-L1 and its PD-associated variants, namely UCH-L1^I93M^ and UCH-L1^S18Y^, is elevated in response to the inhibition of the UPS, which provides a functional link between PD and the UPS [41]. In the following sections, we discuss the current understanding of disease-associated mutations in UCH-L1.

## 2. DISEASE-ASSOCIATED MUTATIONS IN UCH-L1

### 2.1 p.I93M MUTATION IS ASSOCIATED WITH INCREASED RISK OF PD

The autosomal-dominant p.I93M mutation in UCH-L1 (UCH-L1^I93M^) was first identified in a German family with late-onset PD symptoms, but this report was the only case of this particular familial missense mutation that has been documented in the literature [8]. The human UCH-L1^I93M^ transgenic mice also showed behavioral and physiological phenotypes of Parkinsonism, decreased spontaneous and voluntary movement, and dopaminergic neuronal loss in SN at about 20 weeks of age [68]. However, histological studies of *gad* mice show decreased monoUb level and formation of protein inclusion *in vivo*, without further development of Parkinsonism such as cell degeneration in SN [31]. The two results derived from animal models imply the UCH-L1^I93M^-associated neurotoxicity and its role as a risk factor in PD. At the molecular level, UCH-L1^I93M^ features an increased aggregation propensity both in an UCH-L1^I93M^ transgenic mouse model [68, 69] and transfection cell line [41]. The DUB activity is reduced by half as compared with the wild type (Table 2) [8, 43]. Recently, UCH-L1^I93M^ was found to have aberrant molecular interactions with HSP90 and HSC70, involved in CMA [42]. Although UCH-L1 does not contain the conserved CMA-targeting motif KFERQ [70] recognized by HSC70 in its primary sequence, a putative motif RKKQIE [71] that resembles the CMA-targeting motif exists in helix 2 of UCH-L1 but not in UCH-L3. Nevertheless, we purified recombinant HSP90 and HSC70 to test their physical interactions with UCH-L1 *in vitro*, but the results were negative, which suggests that interactions between UCH-L1 and the molecular chaperones may be regulated by additional functions (Jackson SE, Werrel EF and Hsu STD, unpublished data). Because wild-type UCH-L1 also binds LAMP-2A of the CMA machinery, UCH-L1 variants may interfere in CMA to block αS clearance through a chaperone-independent pathway [72].

As mentioned in Section 1.2, HNE-modified UCH-L1 shares similar phenotype with UCH-L1^I93M^ [43], suggesting that UCH-L1^I93M^ might induce progressive neurodegeneration through altered interactions with other associated proteins such as the previously mentioned CMA components or increased population of toxic aggregates of UCH-L1^I93M^ itself. Indeed, UCH-L1^I93M^ shows increased protein aggregation propensity in different mammalian cell lines [41] (COS-7, Neuro2a and SH-SY5Y cells) and in brain tissues of UCH-L1^I93M^ transgenic mice [68, 69]. The p.I93M mutation reduces the Ub hydrolase activity of UCH-L1 by approximately 50% (Table 2), which may be attributed to the conformation perturbations induced by the p.I93M substitution that is adjacent to the catalytic cysteine residue, Cys90, at the catalytic cleft (Figure 3b), but does not perturb monoUb binding affinity. In the same study, far-UV CD spectroscopy demonstrated reduced secondary structure content with p.I93M mutation as compared with the wild type. Nonetheless, superimposing the crystal structures of UCH-L1^WT^ with those of the PD-associated variants, either in apo forms or in complex with ubiquitin vinyl methyl ester (Ub-VME), yielded a positional root mean square deviation (RMSD) of the backbone Cα atoms of < 0.3 Å, so the PD-associated mutations resulted in negligible structural perturbations in the crystalline states (Figure 4a) [40]. Recently, we reported on a comprehensive folding study of the impact of PD-associated mutations on UCH-L1 by solution-state NMR, intrinsic fluorescence and far-UV CD spectroscopy [73]. First, structural mapping of the observed chemical shift perturbations by the p.I93M mutation showed long-range structural perturbations beyond the mutation site (Figure 4c), which were subsequently confirmed side-chain methyl NMR chemical shift perturbations [74]. Second, NMR hydrogen-deuterium exchange (HDX) experiments showed that the observed backbone amide protection factors, which reflect the stability of the hydrogen bonds that prevents solvent exchange with the corresponding amide groups, are significantly reduced as a result of the p.I93M mutation (Figure 5). Finally, the stopped-flow fluorescence analyses showed that UCHL1^I93M^ exhibits accelerated global unfolding kinetics by approximately one order of magnitude compared with the wild type [73]. Therefore could be speculated that while the p.I93M mutation does not lead to significant unfolding of the solution structure of UCH-L1, the reduced folding stability, which marginally increases the partially unfolded intermediate populations. This situation could lead to accumulation of the aggregation-prone species over a long period of time leading eventually to cellular toxicity, with reduced ability to maintain proteostasis, which is consistent with the age-dependent onset of PD.

**Figure 4.**
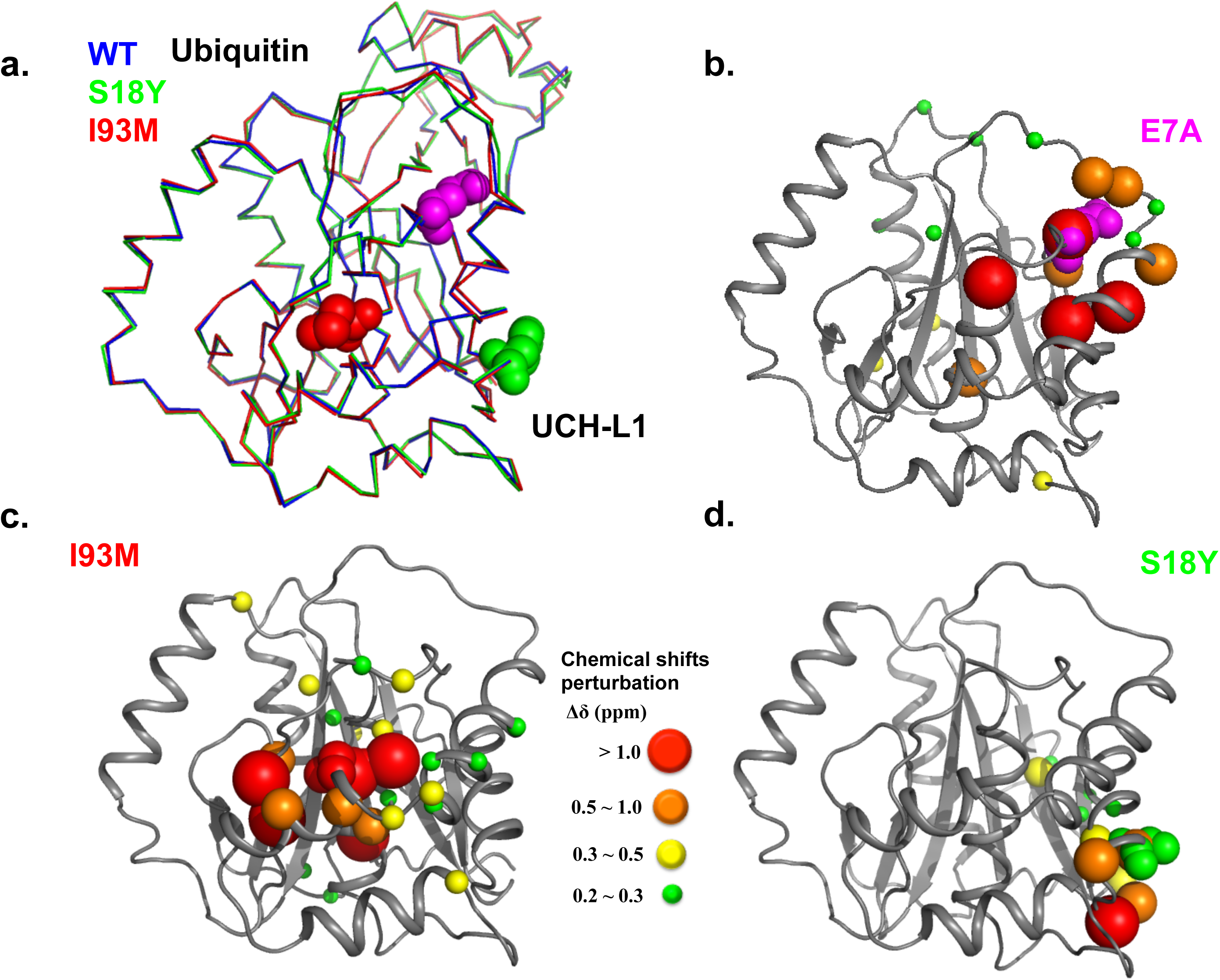
Structural comparison of UCH-L1 variants. **a.** The crystal structures of UCH-L1^I93M^ (dark grey) and UCH-L1^S18Y^ (light grey) are aligned with the structure of a suicide complex of wild-type UCH-L1 in complex Ub-VME (black). Ub is shown in grey surface representation. The mutations associated with neurodegenerative diseases, i.e., p.I93M (**b**), p.S18Y (**c**) and p.E7A (**d**), are shown in sticks. Structural mapping of NMR chemical shift perturbations; the size of chemical shift perturbations are shown in different radii of the backbone amide nitrogen atoms using the crystal structure of wild-type UCH-L1 (PDB ID: 2ETL) as template.

**Figure 5.**
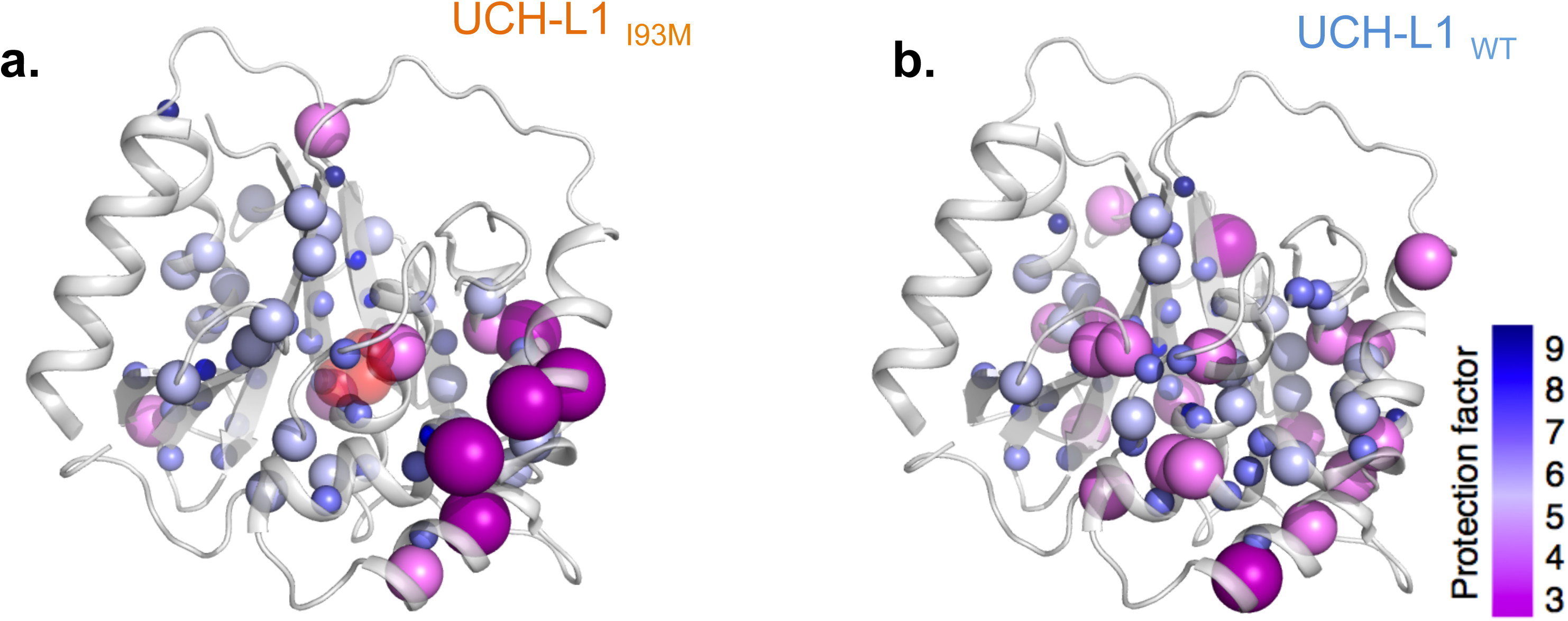
NMR HDX analysis of UCH-L1 variants. The amide nitrogen atoms are shown in spheres and are color-ramped from light grey to white to dark grey according to the scale bar for the NMR HDX-derived protection factors. The radii of the spheres are proportional to the magnitudes of the observed protection factors.

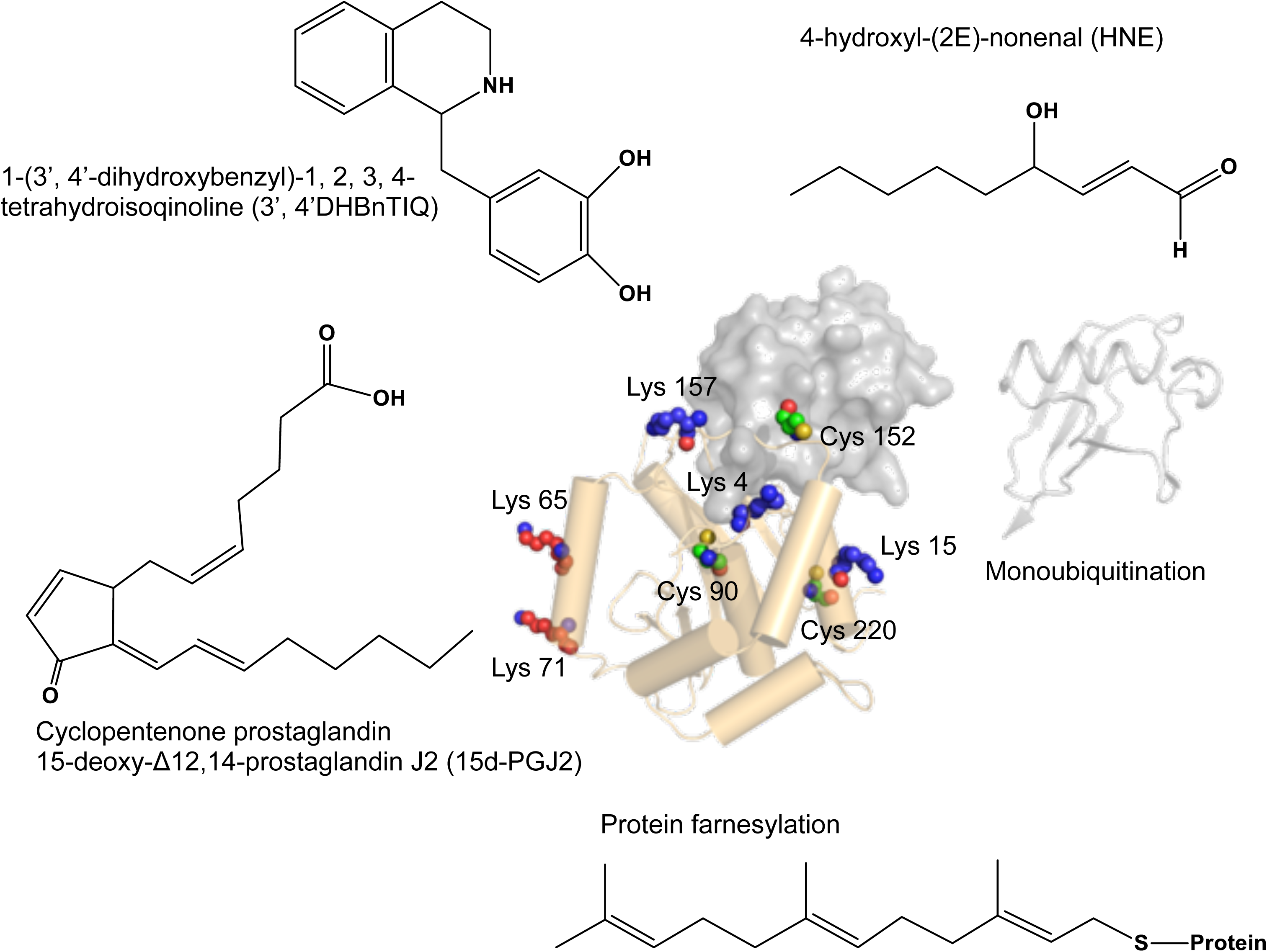

### 2.2. CONTROVERSIAL GENETIC LINK BETWEEN p.S18Y MUTATION AND PD

The p.S18Y amino acid substitution was originally found to play a protective role in early-onset PD for Scandinavians [75]. The protective role of the p.S18Y variant was also reported for Chinese [76], Japanese [77-79], and German populations [80]. Recently, an international consortium has been established to survey the genetic link between the p.S18Y variant of UCH-L1 (UCH-L1^S18Y^) and PD: a similar conclusion about the protective role in PD has been reached [81]. Nevertheless, contradictory results have been reported for populations in Italy [82], Australia [83], European Caucasians in the United States [78], Han Chinese [84, 85], and more recently Japanese [86], which is possibly due to differences in the ethnic population. Therefore, the implication of the p.S18Y variant in PD remains to be determined. Although a number of studies have suggested that UCH-L1^S18Y^ may reduce susceptibility to PD [49, 87], little is known about the molecular mechanism of the neuroprotective functions of UCH-L1^S18Y^. One report linked UCH-L1^S18Y^ to the UPS pathway, showing that although proteasome inhibitor treatment in the COS-7 cell culture significantly increased the aggregation of UCH-L1^I93M^, the number of cellular inclusions could be significantly reduced by co-transfection with UCH-L1^S18Y^, which suggests that UCH-L1^S18Y^ could suppress the cellular toxicity of UCH-L1^I93M^ *in trans* [41]. Despite no precedence of UCH-L1^S18Y/I93M^ compound heterozygosity in humans as has been designed for the artificial cell biology study, the protective role of UCH-L1^S18Y^ could also be manifested by its role in protecting against oxidative stress-related PD pathogenesis [87, 88]. In addition, the previously reported reduced dimerization tendency of UCH-L1^S18Y^ that antagonizes the unusual Ub ligase activity may play a role in conferring the protective role against PD.

### 2.3. p.E7A MUTATION CAUSES AUTOSOMAL-RECESSIVE EARLY-ONSET NEURODEGENERATION

Recently, p.E7A mutation in UCH-L1 (UCH-L1^E7A^) was identified in a Turkish family with autosomal-recessive early-onset of progressive neurodegeneration involving impaired eyesight in childhood, cerebellar ataxia, and spasticity with upper motor-neuron dysfunction [9]. While this disease shares little symptomatic overlap with PD, it does resemble some of the phenotypes of the *gad* mouse, such as ataxia. At the molecular level, UCH-L1^E7A^ shows significant loss of Ub binding capacity thereby leading to nearly complete loss of DUB activity (Table 2). Such a loss of function can be rationalized structurally because Glu7 is highly conserved among all UCHs (Figure 2) and is located at the Ub binding interface to form electrostatic interactions with Arg42 and Arg72 of Ub (Figure 3). One the one hand, the loss of DUB activity of UCH-L1^E7A^ may be related to early-onset neurodegeneration despite the fact that UCH-L1 is not a very effective DUB compared to other DUBs, such as UCH-L3. On the other hand, the loss of Ub binding affinity is the result of the inability of UCH-L1^E7A^ to form stable complex with Ub. Therefore, it is plausible that the loss of monoUb stabilization effect is responsible for this form of neurodegeneration. The exact molecular mechanism underlying the neurodegeneration remains to be established.

## 3. OXIDATIVE DAMAGE AND PTMS OF UCH-L1

### 3.1. MEMBRANE-ASSOCIATED FARNESYLATION OF UCH-L1 PROMOTES ALPHA-SYNUCLEIN NEUROTOXICITY

Membrane-associated UCH-L1 was found in mouse ova and brain cells [89] whose amount promotes intracellular αS induced neurotoxicity [45, 90]. The membrane-associated UCH-L1 can be down-regulated by treatment with the farnesyltransferase inhibitor (FTI-277) and/or by introducing a p.C220S point mutation in UCH-L1, which suggests that the Cys220 site-specific farnesylation of UCH-L1 is responsible for its membrane association. Instead of the conventional sequence of protein farnesylation (CaaX motif consists of a cysteine residue followed by two aliphatic amino acids and an end residue X: S/M/Q/A/T) that was used for membrane association of small GTPase (Ras), UCH-L1 contains an atypical farnesylation sequence at its C-terminus (C_220_KAA-COOH). Coexpression of αS and the UCH-L1 variant that harbors an optimal farnesylated sequence, CIVM, at the C-terminal sequence resulted in increased cellular toxicity in SH-SY5Y human neuroblastoma cells [45]. However, another recent study reported that the amount of membrane-associated UCH-L1 is independent of C-terminal farnesylation using different neuron cell types [90]. Given that inhibition of farnesylation did not alter the sub-localization of membrane-associated UCH-L1 in cultured cortical neurons and in the mouse Neuron2a cell line, the results suggest that UCH-L1 may exhibit intrinsic affinity towards cell membrane or membrane-associated proteins in different cell types. The connection between membrane-association and αS aggregation remains to be established with further experimental evidence. Nevertheless, these results suggest that altering sub-localization of UCH-L1 between membrane and cytosol is linked to αS induced cellular toxicity and could be served as a new therapeutic target of PD.

### 3.2. N-TERMINALLY TRUNCATED UCH-L1 MIGHT PREVENT NEURONS FROM PD-ASSOCIATED DAMAGE

The N-terminally truncated UCH-L1 (UCH-L1^NT^) isoform was first identified in human brain tissue in a proteomic study [91]. UCH-L1_NT_ was subsequently identified in an SH-SY5Y cell line and lung cancer cell line (NCI-H157) and in whole mouse-brain tissue [48]. Mass spectrometry-based hydrogen-deuterium exchange analysis showed that the truncation of the first 11 residues at the N-terminus markedly decreased overall structural stability and DUB activity of UCH-L1 [48]. Interestingly, UCH-L1^NT^ showed protective effects when it was recruited onto the mitochondrial outer membrane. On mitochondrial recruitment, UCH-L1^NT^ self-dimerizes by forming a disulfide bond between the catalytic Cys90 side-chains. A suggestion was made that the self-dimerization could prevent cells from oxidative damage by inhibiting mitochondrial complex I. Such effect, however, was not observed for full-length UCH-L1 under the same oxidative condition. Furthermore, monoubiquitination of UCH-L1^NT^ at position Lys15 or Lys157 could accelerate UCH-L1^NT^ turnover rate, which competes with the UCH-L1^NT^ function that prevents PD-associated oxidative damage.

### 3.3. REVERSIBLE MONO-UBIQUITINATION OF UCH-L1 REGULATES MONO-UBIQUITIN BINDING

Mono-ubiquitination is observed in UCH-L1^NT^ and in overexpressed full-length UCH-L1 in COS-7 cells [46]. Mono-ubiquitination was identified for four conserved lysines in UCH-L1, namely, Lys4, Lys65, Lys71, and Lys157. Additionally, a p.K157R replacement at the crossover loop resulted in markedly reduced mono-ubiquitination level, which suggests that Lys157 may be the preferred ubiquitination site of UCH-L1. Given that UCH-L1 binds to and stabilizes monoUb, the mono-ubiqutination at Lys157 close to the monoUb binding interface is expected to generate significant steric hindrance against substrate recognition of UCHs [38]. Moreover, the catalytically inactive mutations (p.C90S and p.D176N) as well as the mutation (p.D30K) in the monoUb binding interface change the basal level of mono-ubiquitinated UCH-L1, suggesting that the mono-ubiquitination of UCH-L1 could be tightly associated with DUB activity of UCH-L1.

### 3.4. POST-TRANSLATION MODIFICATIONS OF UCH-L1 BY REACTIVE METABOLITES INDUCE UNFOLDING AND AGGREGATION OF UCH-L1

During the inflammatory and oxidative response, various lipid-derived metabolites and reactive oxygen species can modify the side-chains of lysine and cysteine residues, thereby causing protein dysfunction. PTMs of UCH-L1 by these reactive lipid metabolites have been reported [92]. In particular, Gronenborn and co-workers demonstrated that cyclopentenone prostaglandin, 15-deoxy- Δ 12, 14-prostaglandin J2 (15d-PGJ2) specifically reacts with Cys152 on the crossover loop of UCH-L1 [93]. According to hetero-nuclear NMR spectroscopy analysis, 15d-PGJ2–modification resulted in significant NMR line broadening and chemical shift perturbations, thereby leading to partial unfolding and aggregation, which was subsequently confirmed by analytical size-exclusion chromatography. Such PTM resembles the effect of HNE modification (Sections 1.2 ad 2.1). Note that while prostaglandin primarily modifies Cys152 on the crossover loop, the small size of HNE enables its access to protein interiors for side-chain modifications. Systematic mutagenesis indicated that Cys90, in addition to Cys152, is the initial site for HNE modification [92], which is not surprising because Cys90 is the most reactive group of the catalytic triad within UCH-L1. In addition to lipid metabolites, a dopamine derivative 1-(3’,4’-dihydroxybenzyl)-1,2,3,4-tetrahydroisoquinoline (3’,4’ DHBnTIQ) can also modify Cys152 of UCH-L1 and result in similar effects as those induced by 15d-PGJ2 modification [94]. Together, these results indicate that UCH-L1 is highly susceptible to PTMs that can lead to partial unfolding and increased aggregation propensity resulting in similar phenotypes as that of UCH-L1^I93M^ (Section 2.1).

## 4. DISCUSSION

DUBs are involved in multiple biological functions, including malignancy and neurodegenerative diseases [16, 95]. UCH-L1 is a highly expressed neuro-specific DUB that has been linked to PD, but little is known about its specific functions or the molecular machineries that participate in cellular regulations. Because UCH-L1 cannot process isopeptide bonds within polyUb chains and possesses low hydrolase activity as compared with other DUBs, loss of UCH-L1 DUB activity due to mutations or PTMs could be compensated by other DUBs and may not significantly perturb the protein homeostasis involving the UPS. In other words, loss of DUB function is unlikely a major contributor to the UCH-L1 effect in the context of neurodegenerative diseases.

Although emerging reports have suggested that UCH-L1 has important functions independent of its DUB activity, including Ub ligase activity, membrane association, subcellular localization, interactions with CDKs and aberrant interactions with CMA components, the molecular details of these interactions are lacking. Because a PD-associated variant such as UCH-L1^I93M^ displays increased aggregation propensity both *in vivo* and *in vitro*, the impact of PD-related mutations and endogenous reactive molecule modifications on UCH-L1 may be rationalized as a misfolding-related disorder. Indeed, the familial p.I93M mutation, HNE modification and oxidation to catalytic Cys90, which resides in helix 3, and cyclopentenone prostaglandin modification to Cys152 on the crossover loop decrease the global protein stability and dynamics of UCH-L1. These modifications are located near the catalytic site or at the Ub binding interface. However, the p.S18Y mutation at the surface of UCH-L1 does not significantly perturb the structural stability of UCH-L1 but reduces the suggested function of Ub ligase relying on UCH-L1 dimerization. Reduction of UCH-L1 ligase activity affects the clearance of Lys63 linkage-specific polyubiquitination of 〈S?, which might be the major mechanistic part involved in PD. p.E7A mutation is a high risk factor in the autosomal-recessive early-onset neurodegenerative disorder and also a highly conserved residue in the monoUb-binding interface (Figure 3). Reducing monoUb binding is the primary effect of UCH-L1^E7A^ and resembles that of the p.D30K mutation at the Ub binding interface. In contrast to the wealth of biochemical insights, much less is understood structurally in terms of the impact of p.E7A mutation in neurodegenerative diseases. In light of the close genetic link between UCH-L1^E7A^ and neurodegenerative disorders, further insights into how the p.E7A mutation affects neurological functions are necessary.

The proposed function of UCH-L1 as a monoUb stabilizer [35] deserves further investigation given that Ub is one of the most stable proteins, it can withstand very harsh physico-chemical conditions and it is involved in direct interactions with the proteasome for protein degradation. The molecular basis of such a stabilization effect remains elusive because UCH-L1 is chemically and thermally much less stable than Ub [73]. Nevertheless, UCH-L1 binds to Ub with very high affinity, slow-off rate [37] and an extensive binding interface. The sequestered solvent accessible surface area is 2732 Å^2^ according to the crystal structure of UCH-L1 in complex with Ub (PDB entry: 3kw5). Given the cellular abundance of the molecules, the mechanism by which Ub is released from UCH-L1 to carry out its physiological functions along the UPS and signaling pathways in neurons remains elusive. Understanding the importance of Ub binding independent of its DUB activity may also shed light on the underlying mechanism of neurodegeneration associated with the p.E7A mutation. Finally, little is known about the proportion of post-translationally modified UCH-L1 by HNE, 3’,4’DHBnTIQ, cyclopentenone prostaglandin, and ubiquitination under physiological conditions. Although *in vitro* and cellular characterizations have established that most of these PTMs can lead to misfolding and aggregation as well as perturbation of important cellular machineries such as CMA [28, 42, 96], whether these aberrant interactions, which also resemble those of the p.I93M familial mutation, actually lead to gain of toxicity associated with Parkinsonism remains to be addressed in molecular detail.

## ACKNOWLEDGEMENT

STDH is a recipient of a Career Development Award of the International Human Frontier Science Program (CDA-00025/2010-C). This work is in part supported by the Ministry of Science and Technology (100-2113-M-001-031-MY2 and 102-2113-M-001-017-MY2), National Tsing Hua University, the National Synchrotron Radiation Research Center, and Academia Sinica, Taiwan.

## 5. REFERENCES

[1] Chung, S.J.; Armasu, S.M.; Biernacka, J.M.; Lesnick, T.G.; Rider, D.N.; Lincoln, S.J.; Ortolaza, A.I.; Farrer, M.J.; Cunningham, J.M.; Rocca, W.A., et al. Common variants in park loci and related genes and Parkinson’s disease. Mov Disord, 2011. 26(2), 280-8.

[2] Lesage, S.; Brice, A. Parkinson’s disease: From monogenic forms to genetic susceptibility factors. Hum Mol Genet, 2009. 18(R1), R48-59.

[3] Meriin, A.B.; Sherman, M.Y. Role of molecular chaperones in neurodegenerative disorders. Int J Hyperthermia, 2005. 21(5), 403-419.

[4] Castro-Caldas, M.; Carvalho, A.N.; Rodrigues, E.; Henderson, C.J.; Wolf, C.R.; Rodrigues, C.M.; Gama, M.J. Tauroursodeoxycholic acid prevents MPTP-induced dopaminergic cell death in a mouse model of Parkinson’s disease. Mol Neurobiol, 2012. 46(2), 475-486.

[5] Xia, L.P.; Li, L.Y.; Fei, X.F.; Liang, Z.Q. Autophagy is involved in 6-OHDA-induced dopaminergic cell death. Nan Fang Yi Ke Da Xue Xue Bao, 2010. 30(12), 2649-2651.

[6] Radad, K.; Rausch, W.D.; Gille, G. Rotenone induces cell death in primary dopaminergic culture by increasing ros production and inhibiting mitochondrial respiration. Neurochem Int, 2006. 49(4), 379-386.

[7] Shavali, S.; Ebadi, M. 1-benzyl-1,2,3,4-tetrahydroisoquinoline (1BnTIQ), an endogenous neurotoxin, induces dopaminergic cell death through apoptosis. Neurotoxicology, 2003. 24(3), 417-424.

[8] Leroy, E.; Boyer, R.; Auburger, G.; Leube, B.; Ulm, G.; Mezey, E.; Harta, G.; Brownstein, M.J.; Jonnalagada, S.; Chernova, T., et al. The ubiquitin pathway in Parkinson’s disease. Nature, 1998. 395(6701), 451-452.

[9] Bilguvar, K.; Tyagi, N.K.; Ozkara, C.; Tuysuz, B.; Bakircioglu, M.; Choi, M.; Delil, S.; Caglayan, A.O.; Baranoski, J.F.; Erturk, O., et al. Recessive loss of function of the neuronal ubiquitin hydrolase UCHL1 leads to early-onset progressive neurodegeneration. Proc Natl Acad Sci U S A, 2013. 110(9), 3489-3494.

[10] Setsuie, R.; Wada, K. The functions of UCH-L1 and its relation to neurodegenerative diseases. Neurochem Int, 2007. 51(2-4), 105-111.

[11] Achey, M.; Aldred, J.L.; Aljehani, N.; Bloem, B.R.; Biglan, K.M.; Chan, P.; Cubo, E.; Dorsey, E.R.; Goetz, C.G.; Guttman, M., et al. The past, present, and future of telemedicine for Parkinson’s disease. Mov Disord, 2014. 29(7), 871-883.

[12] Thompson, R.J.; Doran, J.F.; Jackson, P.; Dhillon, A.P.; Rode, J. PGP 9.5--a new marker for vertebrate neurons and neuroendocrine cells. Brain Res, 1983. 278(1-2), 224-228.

[13] Day, I.N. Enolases and PGP9.5 as tissue-specific markers. Biochem Soc Trans, 1992. 20(3), 637-642.

[14] Day, I.N.; Thompson, R.J. UCHL1 (PGP 9.5): Neuronal biomarker and ubiquitin system protein. Prog Neurobiol, 2010. 90(3), 327-362.

[15] Komander, D. Mechanism, specificity and structure of the deubiquitinases. Subcell Biochem, 2010. 54, 69-87.

[16] Hussain, S.; Zhang, Y.; Galardy, P.J. Dubs and cancer: The role of deubiquitinating enzymes as oncogenes, non-oncogenes and tumor suppressors. Cell Cycle, 2009. 8(11), 1688-1697.

[17] Fang, Y.; Fu, D.; Shen, X.Z. The potential role of ubiquitin c-terminal hydrolases in oncogenesis. Biochim Biophys Acta, 2010. 1806(1), 1-6.

[18] Jara, J.H.; Frank, D.D.; Ozdinler, P.H. Could dysregulation of ups be a common underlying mechanism for cancer and neurodegeneration? Lessons from UCHL1. Cell Biochem Biophys, 2013. 67(1), 45-53.

[19] Ma, Y.; Zhao, M.; Zhong, J.; Shi, L.; Luo, Q.; Liu, J.; Wang, J.; Yuan, X.; Huang, C. Proteomic profiling of proteins associated with lymph node metastasis in colorectal cancer. J Cell Biochem, 2010. 110(6), 1512-1519.

[20] Schroder, C.; Milde-Langosch, K.; Gebauer, F.; Schmid, K.; Mueller, V.; Wirtz, R.M.; Meyer-Schwesinger, C.; Schluter, H.; Sauter, G.; Schumacher, U. Prognostic relevance of ubiquitin C-terminal hydrolase L1 (UCH-L1) mRNA and protein expression in breast cancer patients. J Cancer Res Clin Oncol, 2013. 139(10), 1745-1755.

[21] Hussain, S.; Foreman, O.; Perkins, S.L.; Witzig, T.E.; Miles, R.R.; van Deursen, J.; Galardy, P.J. The de-ubiquitinase UCH-L1 is an oncogene that drives the development of lymphoma in vivo by deregulating phlpp1 and Akt signaling. Leukemia, 2010. 24(9), 1641-1655.

[22] Bheda, A.; Yue, W.; Gullapalli, A.; Shackelford, J.; Pagano, J.S. PU.1-dependent regulation of UCH L1 expression in b-lymphoma cells. Leuk Lymphoma, 2011. 52(7), 1336-1347.

[23] Jang, M.J.; Baek, S.H.; Kim, J.H. UCH-L1 promotes cancer metastasis in prostate cancer cells through emt induction. Cancer Lett, 2011. 302(2), 128-135.

[24] Orr, K.S.; Shi, Z.; Brown, W.M.; O’Hagan, K.A.; Lappin, T.R.; Maxwell, P.; Percy, M.J. Potential prognostic marker ubiquitin carboxyl-terminal hydrolase-L1 does not predict patient survival in non-small cell lung carcinoma. J Exp Clin Cancer Res, 2011. 30, 79.

[25] Liu, Y.; Lashuel, H.A.; Choi, S.; Xing, X.; Case, A.; Ni, J.; Yeh, L.A.; Cuny, G.D.; Stein, R.L.; Lansbury, P.T., Jr. Discovery of inhibitors that elucidate the role of UCH-L1 activity in the H1299 lung cancer cell line. Chem Biol, 2003. 10(9), 837-846.

[26] Goto, Y.; Zeng, L.; Yeom, C.J.; Zhu, Y.; Morinibu, A.; Shinomiya, K.; Kobayashi, M.; Hirota, K.; Itasaka, S.; Yoshimura, M., et al. UCHL1 provides diagnostic and antimetastatic strategies due to its deubiquitinating effect on HIF-1α. Nat Commun, 2015. 6, 6153.

[27] Frisan, T.; Coppotelli, G.; Dryselius, R.; Masucci, M.G. Ubiquitin c-terminal hydrolase-L1 interacts with adhesion complexes and promotes cell migration, survival, and anchorage independent growth. FASEB J, 2012. 26(12), 5060-5070.

[28] Kabuta, T.; Mitsui, T.; Takahashi, M.; Fujiwara, Y.; Kabuta, C.; Konya, C.; Tsuchiya, Y.; Hatanaka, Y.; Uchida, K.; Hohjoh, H., et al. Ubiquitin C-terminal hydrolase L1 (UCH-L1) acts as a novel potentiator of cyclin-dependent kinases to enhance cell proliferation independently of its hydrolase activity. J Biol Chem, 2013. 288(18), 12615-12626.

[29] Ummanni, R.; Jost, E.; Braig, M.; Lohmann, F.; Mundt, F.; Barett, C.; Schlomm, T.; Sauter, G.; Senff, T.; Bokemeyer, C., et al. Ubiquitin carboxyl-terminal hydrolase 1 (UCHL1) is a potential tumour suppressor in prostate cancer and is frequently silenced by promoter methylation. Mol. Cancer, 2011. 10.

[30] Bittencourt Rosas, S.L.; Caballero, O.L.; Dong, S.M.; da Costa Carvalho Mda, G.; Sidransky, D.; Jen, J. Methylation status in the promoter region of the human PGP9.5 gene in cancer and normal tissues. Cancer Lett, 2001. 170(1), 73-79.

[31] Saigoh, K.; Wang, Y.L.; Suh, J.G.; Yamanishi, T.; Sakai, Y.; Kiyosawa, H.; Harada, T.; Ichihara, N.; Wakana, S.; Kikuchi, T., et al. Intragenic deletion in the gene encoding ubiquitin carboxy-terminal hydrolase in gad mice. Nat Genet, 1999. 23(1), 47-51.

[32] Chen, F.J.; Sugiura, Y.; Myers, K.G.; Liu, Y.; Lin, W.C. Ubiquitin carboxyl-terminal hydrolase L1 is required for maintaining the structure and function of the neuromuscular junction. Proc. Natl. Acad. Sci. U.S.A., 2010. 107(4), 1636-1641.

[33] Kurihara, L.J.; Kikuchi, T.; Wada, K.; Tilghman, S.M. Loss of UCH-L1 and UCH-L3 leads to neurodegeneration, posterior paralysis and dysphagia. Hum Mol Genet, 2001. 10(18), 1963-1970.

[34] Shimshek, D.R.; Schweizer, T.; Schmid, P.; van der Putten, P.H. Excess alpha-synuclein worsens disease in mice lacking ubiquitin carboxy-terminal hydrolase L1. Sci Rep, 2012. 2, 262.

[35] Osaka, H.; Wang, Y.L.; Takada, K.; Takizawa, S.; Setsuie, R.; Li, H.; Sato, Y.; Nishikawa, K.; Sun, Y.J.; Sakurai, M., et al. Ubiquitin carboxy-terminal hydrolase L1 binds to and stabilizes monoubiquitin in neuron. Hum Mol Genet, 2003. 12(16), 1945-1958.

[36] Cartier, A.E.; Djakovic, S.N.; Salehi, A.; Wilson, S.M.; Masliah, E.; Patrick, G.N. Regulation of synaptic structure by ubiquitin C-terminal hydrolase L1. J. Neurosci., 2009. 29(24), 7857-7868.

[37] Case, A.; Stein, R.L. Mechanistic studies of ubiquitin C-terminal hydrolase L1. Biochemistry, 2006. 45(7), 2443-2452.

[38] Zhou, Z.R.; Zhang, Y.H.; Liu, S.; Song, A.X.; Hu, H.Y. Length of the active-site crossover loop defines the substrate specificity of ubiquitin c-terminal hydrolases for ubiquitin chains. Biochem J, 2012. 441(1), 143-149.

[39] Larsen, C.N.; Krantz, B.A.; Wilkinson, K.D. Substrate specificity of deubiquitinating enzymes: Ubiquitin c-terminal hydrolases. Biochemistry, 1998. 37(10), 3358-68.

[40] Boudreaux, D.A.; Maiti, T.K.; Davies, C.W.; Das, C. Ubiquitin vinyl methyl ester binding orients the misaligned active site of the ubiquitin hydrolase UCHL1 into productive conformation. Proc Natl Acad Sci U S A, 2010. 107(20), 9117-9122.

[41] Ardley, H.C.; Scott, G.B.; Rose, S.A.; Tan, N.G.; Robinson, P.A. UCH-L1 aggresome formation in response to proteasome impairment indicates a role in inclusion formation in Parkinson’s disease. J Neurochem, 2004. 90(2), 379-391.

[42] Kabuta, T.; Furuta, A.; Aoki, S.; Furuta, K.; Wada, K. Aberrant interaction between Parkinson disease-associated mutant UCH-L1 and the lysosomal receptor for chaperone-mediated autophagy. J Biol Chem, 2008. 283(35), 23731-23738.

[43] Nishikawa, K.; Li, H.; Kawamura, R.; Osaka, H.; Wang, Y.L.; Hara, Y.; Hirokawa, T.; Manago, Y.; Amano, T.; Noda, M., et al. Alterations of structure and hydrolase activity of Parkinsonism-associated human ubiquitin carboxyl-terminal hydrolase L1 variants. Biochem Biophys Res Commun, 2003. 304(1), 176-183.

[44] Cole, R.N.; Hart, G.W. Cytosolic O-glycosylation is abundant in nerve terminals. J Neurochem, 2001. 79(5), 1080-1089.

[45] Liu, Z.; Meray, R.K.; Grammatopoulos, T.N.; Fredenburg, R.A.; Cookson, M.R.; Liu, Y.; Logan, T.; Lansbury, P.T., Jr. Membrane-associated farnesylated UCH-L1 promotes alpha-synuclein neurotoxicity and is a therapeutic target for Parkinson’s disease. Proc Natl Acad Sci U S A, 2009. 106(12), 4635-4640.

[46] Meray, R.K.; Lansbury, P.T., Jr. Reversible monoubiquitination regulates the Parkinson disease-associated ubiquitin hydrolase UCH-L1. J Biol Chem, 2007. 282(14), 10567-10575.

[47] Paige, J.S.; Xu, G.; Stancevic, B.; Jaffrey, S.R. Nitrosothiol reactivity profiling identifies s-nitrosylated proteins with unexpected stability. Chem Biol, 2008. 15(12), 1307-1316.

[48] Kim, H.J.; Kim, H.J.; Jeong, J.E.; Baek, J.Y.; Jeong, J.; Kim, S.; Kim, Y.M.; Kim, Y.; Nam, J.H.; Huh, S.H., et al. N-terminal truncated UCH-L1 prevents Parkinson’s disease associated damage. PLoS ONE, 2014. 9(6), e99654.

[49] Liu, Y.; Fallon, L.; Lashuel, H.A.; Liu, Z.; Lansbury, P.T., Jr. The UCH-L1 gene encodes two opposing enzymatic activities that affect alpha-synuclein degradation and Parkinson’s disease susceptibility. Cell, 2002. 111(2), 209-218.

[50] Bheda, A.; Gullapalli, A.; Caplow, M.; Pagano, J.S.; Shackelford, J. Ubiquitin editing enzyme uch L1 and microtubule dynamics: Implication in mitosis. Cell Cycle, 2010. 9(5), 980-994.

[51] Naito, S.; Mochizuki, H.; Yasuda, T.; Mizuno, Y.; Furusaka, M.; Ikeda, S.; Adachi, T.; Shimizu, H.M.; Suzuki, J.; Fujiwara, S., et al. Characterization of multimetric variants of ubiquitin carboxyl-terminal hydrolase L1 in water by small-angle neutron scattering. Biochem Biophys Res Commun, 2006. 339(2), 717-725.

[52] Nishio, K.; Kim, S.W.; Kawai, K.; Mizushima, T.; Yamane, T.; Hamazaki, J.; Murata, S.; Tanaka, K.; Morimoto, Y. Crystal structure of the de-ubiquitinating enzyme UCH37 (human UCH-L5) catalytic domain. Biochem Biophys Res Commun, 2009. 390(3), 855-860.

[53] Dikic, I.; Wakatsuki, S.; Walters, K.J. Ubiquitin-binding domains - from structures to functions. Nat Rev Mol Cell Biol, 2009. 10(10), 659-671.

[54] Dang, L.C.; Melandri, F.D.; Stein, R.L. Kinetic and mechanistic studies on the hydrolysis of ubiquitin C-terminal 7-amido-4-methylcoumarin by deubiquitinating enzymes. Biochemistry, 1998. 37(7), 1868-1879.

[55] Boudreaux, D.A.; Chaney, J.; Maiti, T.K.; Das, C. Contribution of active site glutamine to rate enhancement in ubiquitin c-terminal hydrolases. FEBS J, 2012. 279(6), 1106-1118.

[56] Maiti, T.K.; Permaul, M.; Boudreaux, D.A.; Mahanic, C.; Mauney, S.; Das, C. Crystal structure of the catalytic domain of uchl5, a proteasome-associated human deubiquitinating enzyme, reveals an unproductive form of the enzyme. FEBS J, 2011. 278(24), 4917-4926.

[57] Mitsui, T.; Hirayama, K.; Aoki, S.; Nishikawa, K.; Uchida, K.; Matsumoto, T.; Kabuta, T.; Wada, K. Identification of a novel chemical potentiator and inhibitors of UCH-L1 by in silico drug screening. Neurochem Int, 2010. 56(5), 679-686.

[58] Goldenberg, S.J.; McDermott, J.L.; Butt, T.R.; Mattern, M.R.; Nicholson, B. Strategies for the identification of novel inhibitors of deubiquitinating enzymes. Biochem Soc Trans, 2008. 36(Pt 5), 828-832.

[59] Bavikar, S.N.; Spasser, L.; Haj-Yahya, M.; Karthikeyan, S.V.; Moyal, T.; Kumar, K.S.; Brik, A. Chemical synthesis of ubiquitinated peptides with varying lengths and types of ubiquitin chains to explore the activity of deubiquitinases. Angew Chem Int Ed Engl, 2012. 51(3), 758-763.

[60] Ohayon, S.; Spasser, L.; Aharoni, A.; Brik, A. Targeting deubiquitinases enabled by chemical synthesis of proteins. J Am Chem Soc, 2012. 134(6), 3281-3289.

[61] Gong, B.; Leznik, E. The role of ubiquitin c-terminal hydrolase L1 in neurodegenerative disorders. Drug News Perspect, 2007. 20(6), 365-370.

[62] Sultana, R.; Boyd-Kimball, D.; Cai, J.; Pierce, W.M.; Klein, J.B.; Merchant, M.; Butterfield, D.A. Proteomics analysis of the alzheimer’s disease hippocampal proteome. J Alzheimers Dis, 2007. 11(2), 153-164.

[63] Hattori, N.; Mizuno, Y. Pathogenetic mechanisms of parkin in Parkinson’s disease. Lancet, 2004. 364(9435), 722-4.

[64] Gosal, D.; Ross, O.A.; Toft, M. Parkinson’s disease: The genetics of a heterogeneous disorder. Eur J Neurol, 2006. 13(6), 616-627.

[65] Mizuno, Y.; Hattori, N.; Yoshino, H.; Hatano, Y.; Satoh, K.; Tomiyama, H.; Li, Y. Progress in familial Parkinson’s disease. J Neural Transm Suppl, 2006(70), 191-204.

[66] Lowe, J.; McDermott, H.; Landon, M.; Mayer, R.J.; Wilkinson, K.D. Ubiquitin carboxyl-terminal hydrolase (PGP 9.5) is selectively present in ubiquitinated inclusion bodies characteristic of human neurodegenerative diseases. J Pathol, 1990. 161(2), 153-160.

[67] Coleman, M. Axon degeneration mechanisms: Commonality amid diversity. Nat Rev Neurosci, 2005. 6(11), 889-98.

[68] Setsuie, R.; Wang, Y.L.; Mochizuki, H.; Osaka, H.; Hayakawa, H.; Ichihara, N.; Li, H.; Furuta, A.; Sano, Y.; Sun, Y.J., et al. Dopaminergic neuronal loss in transgenic mice expressing the Parkinson’s disease-associated UCH-L1 I93M mutant. Neurochem Int, 2007. 50(1), 119-129.

[69] Yasuda, T.; Nihira, T.; Ren, Y.R.; Cao, X.Q.; Wada, K.; Setsuie, R.; Kabuta, T.; Wada, K.; Hattori, N.; Mizuno, Y., et al. Effects of UCH-L1 on alpha-synuclein over-expression mouse model of Parkinson’s disease. J Neurochem, 2009. 108(4), 932-944.

[70] Cuervo, A.M.; Dice, J.F. Unique properties of LAPM2A compared to other LAMP2 isoforms. J Cell Sci, 2000. 113 Pt 24, 4441-4450.

[71] Bandyopadhyay, U.; Kaushik, S.; Varticovski, L.; Cuervo, A.M. The chaperone-mediated autophagy receptor organizes in dynamic protein complexes at the lysosomal membrane. Mol Cell Biol, 2008. 28(18), 5747-5763.

[72] Cuervo, A.M.; Wong, E. Chaperone-mediated autophagy: Roles in disease and aging. Cell Res, 2014. 24(1), 92-104.

[73] Andersson, F.I.; Werrell, E.F.; McMorran, L.; Crone, W.J.; Das, C.; Hsu, S.-T.D.; Jackson, S.E. The effect of Parkinson’s-disease-associated mutations on the deubiquitinating enzyme UCH-L1. J Mol Biol, 2011. 407(2), 261-272.

[74] Kumar, S.M.; Lyu, P.C.; Hsu, S.-T.D. Structural perturbation of the Parkinson’s disease-associated I93M mutation in human UCH-L1 revealed by solution state NMR spectroscopy. Chinese Journal of Magnetic Resonance, 2015, in press.

[75] Carmine Belin, A.; Westerlund, M.; Bergman, O.; Nissbrandt, H.; Lind, C.; Sydow, O.; Galter, D. S18Y in ubiquitin carboxy-terminal hydrolase L1 (UCH-L1) associated with decreased risk of Parkinson’s disease in Sweden. Parkinsonism Relat Disord, 2007. 13(5), 295-298.

[76] Tan, E.K.; Puong, K.Y.; Fook-Chong, S.; Chua, E.; Shen, H.; Yuen, Y.; Pavanni, R.; Wong, M.C.; Puvan, K.; Zhao, Y. Case-control study of UCHL1 S18Y variant in Parkinson’s disease. Mov Disord, 2006. 21(10), 1765-1768.

[77] Satoh, J.; Kuroda, Y. A polymorphic variation of serine to tyrosine at codon 18 in the ubiquitin c-terminal hydrolase-L1 gene is associated with a reduced risk of sporadic Parkinson’s disease in a japanese population. J Neurol Sci, 2001. 189(1-2), 113-117.

[78] Zhang, J.; Hattori, N.; Leroy, E.; Morris, H.R.; Kubo, S.-I.; Kobayashi, T.; Wood, N.W.; Polymeropoulos, M.H.; Mizuno, Y. Association between a polymorphism of ubiquitin carboxy-terminal hydrolase L1 (UCH-L1) gene and sporadic Parkinson’s disease. Parkinsonism Relat Disord, 2000. 6, 195-197.

[79] Momose, Y.; Murata, M.; Kobayashi, K.; Tachikawa, M.; Nakabayashi, Y.; Kanazawa, I.; Toda, T. Association studies of multiple candidate genes for Parkinson’s disease using single nucleotide polymorphisms. Ann Neurol, 2002. 51(1), 133-136.

[80] Wintermeyer, P.; Kruger, R.; Kuhn, W.; Muller, T.; Woitalla, D.; Berg, D.; Becker, G.; Leroy, E.; Polymeropoulos, M.; Berger, K., et al. Mutation analysis and association studies of the UCHL1 gene in German Parkinson’s disease patients. Neuroreport, 2000. 11(10), 2079-2082.

[81] Maraganore, D.M.; Lesnick, T.G.; Elbaz, A.; Chartier-Harlin, M.C.; Gasser, T.; Kruger, R.; Hattori, N.; Mellick, G.D.; Quattrone, A.; Satoh, J., et al. UCHL1 is a Parkinson’s disease susceptibility gene. Ann Neurol, 2004. 55(4), 512-521.

[82] Savettieri, G.; De Marco, E.V.; Civitelli, D.; Salemi, G.; Nicoletti, G.; Annesi, G.; Ciro Candiano, I.C.; Quattrone, A. Lack of association between ubiquitin carboxy-terminal hydrolase L1 gene polymorphism and PD. Neurology, 2001. 57(3), 560-561.

[83] Mellick, G.D.; Silburn, P.A. The ubiquitin carboxy-terminal hydrolase-L1 gene S18Y polymorphism does not confer protection against idiopathic Parkinson’s disease. Neurosci Lett, 2000. 293(2), 127-130.

[84] Zhang, Z.J.; Burgunder, J.M.; An, X.K.; Wu, Y.; Chen, W.J.; Zhang, J.H.; Wang, Y.C.; Xu, Y.M.; Gou, Y.R.; Yuan, G.G., et al. Lack of evidence for association of a UCH-L1 S18Y polymorphism with Parkinson’s disease in a Han-chinese population. Neurosci Lett, 2008. 442(3), 200-202.

[85] Tan, E.K.; Lu, C.S.; Peng, R.; Teo, Y.Y.; Wu-Chou, Y.H.; Chen, R.S.; Weng, Y.H.; Chen, C.M.; Fung, H.C.; Tan, L.C., et al. Analysis of the UCHL1 genetic variant in Parkinson’s disease among chinese. Neurobiol Aging, 2010. 31(12), 2194-2196.

[86] Miyake, Y.; Tanaka, K.; Fukushima, W.; Kiyohara, C.; Sasaki, S.; Tsuboi, Y.; Yamada, T.; Oeda, T.; Shimada, H.; Kawamura, N., et al. UCHL1 S18Y variant is a risk factor for Parkinson’s disease in Japan. BMC Neurol, 2012. 12, 62.

[87] Kyratzi, E.; Pavlaki, M.; Stefanis, L. The S18Y polymorphic variant of UCH-L1 confers an antioxidant function to neuronal cells. Hum Mol Genet, 2008. 17(14), 2160-71.

[88] Xilouri, M.; Kyratzi, E.; Pitychoutis, P.M.; Papadopoulou-Daifoti, Z.; Perier, C.; Vila, M.; Maniati, M.; Ulusoy, A.; Kirik, D.; Park, D.S., et al. Selective neuroprotective effects of the S18Y polymorphic variant of UCH-L1 in the dopaminergic system. Hum Mol Genet, 2012. 21(4), 874-889.

[89] Sekiguchi, S.; Kwon, J.; Yoshida, E.; Hamasaki, H.; Ichinose, S.; Hideshima, M.; Kuraoka, M.; Takahashi, A.; Ishii, Y.; Kyuwa, S., et al. Localization of ubiquitin C-terminal hydrolase L1 in mouse ova and its function in the plasma membrane to block polyspermy. Am J Pathol, 2006. 169(5), 1722-1729.

[90] Bishop, P.; Rubin, P.; Thomson, A.R.; Rocca, D.; Henley, J.M. UCH-L1 C-terminus plays a key role in protein stability but its farnesylation is not required for membrane association in primary neurons. J Biol Chem, 2014. 289(52), 36140-36149

[91] Choi, J.; Levey, A.I.; Weintraub, S.T.; Rees, H.D.; Gearing, M.; Chin, L.S.; Li, L. Oxidative modifications and down-regulation of ubiquitin carboxyl-terminal hydrolase L1 associated with idiopathic Parkinson’s and alzheimer’s diseases. J Biol Chem, 2004. 279(13), 13256-13264.

[92] Kabuta, T.; Setsuie, R.; Mitsui, T.; Kinugawa, A.; Sakurai, M.; Aoki, S.; Uchida, K.; Wada, K. Aberrant molecular properties shared by familial Parkinson’s disease-associated mutant UCH-L1 and carbonyl-modified UCH-L1. Hum Mol Genet, 2008. 17(10), 1482-1496.

[93] Koharudin, L.M.; Liu, H.; Di Maio, R.; Kodali, R.B.; Graham, S.H.; Gronenborn, A.M. Cyclopentenone prostaglandin-induced unfolding and aggregation of the Parkinson disease-associated UCH-L1. Proc Natl Acad Sci U S A, 2010. 107(15), 6835-6840.

[94] Contu, V.R.; Kotake, Y.; Toyama, T.; Okuda, K.; Miyara, M.; Sakamoto, S.; Samizo, S.; Sanoh, S.; Kumagai, Y.; Ohta, S. Endogenous neurotoxic dopamine derivative covalently binds to Parkinson’s disease-associated ubiquitin c-terminal hydrolase L1 and alters its structure and function. J Neurochem, 2014. 130(6), 826-838.

[95] Ristic, G.; Tsou, W.L.; Todi, S.V. An optimal ubiquitin-proteasome pathway in the nervous system: The role of deubiquitinating enzymes. Front Mol Neurosci, 2014. 7, 72.

[96] Kabuta, T.; Wada, K. Insights into links between familial and sporadic Parkinson’s disease: Physical relationship between UCH-L1 variants and chaperone-mediated autophagy. Autophagy, 2008. 4(6), 827-829.

[97] Nuytemans, K.; Theuns, J.; Cruts, M.; Van Broeckhoven, C. Genetic etiology of Parkinson disease associated with mutations in the SNCA, PARK2, PINK1, PARK7, and LRRK2 genes: A mutation update. Hum Mutat, 2010. 31(7), 763-780.

[98] Kawajiri, S.; Saiki, S.; Sato, S.; Hattori, N. Genetic mutations and functions of PINK1. Trends Pharmacol Sci, 2011. 32(10), 573-580.

[99] Mata, I.F.; Wedemeyer, W.J.; Farrer, M.J.; Taylor, J.P.; Gallo, K.A. Lrrk2 in Parkinson’s disease: Protein domains and functional insights. Trends Neurosci, 2006. 29(5), 286-293.

[100] Ariga, H.; Takahashi-Niki, K.; Kato, I.; Maita, H.; Niki, T.; Iguchi-Ariga, S.M. Neuroprotective function of DJ-1 in Parkinson’s disease. Oxid Med Cell Longev, 2013. 2013, 683-920.

[101] Polymeropoulos, M.H.; Lavedan, C.; Leroy, E.; Ide, S.E.; Dehejia, A.; Dutra, A.; Pike, B.; Root, H.; Rubenstein, J.; Boyer, R., et al. Mutation in the α-synuclein gene identified in families with Parkinson’s disease. Science, 1997. 276(5321), 2045-2047.

[102] Ramirez, A.; Heimbach, A.; Gruendemann, J.; Stiller, B.; Hampshire, D.; Cid, L.P.; Goebel, I.; Mubaidin, A.F.; Wriekat, A.L.; Roeper, J., et al. Hereditary Parkinsonism with dementia is caused by mutations in ATP13A2, encoding a lysosomal type 5 P-type ATPase. Nat. Genet., 2006. 38(10), 1184-1191.

[103] Aharon-Peretz, J.; Badarny, S.; Rosenbaum, H.; Gershoni-Baruch, R. Mutations in the glucocerebrosidase gene and Parkinson disease: Phenotype-genotype correlation. Neurology, 2005. 65(9), 1460-1461.

[104] Pan, T.; Kondo, S.; Le, W.; Jankovic, J. The role of autophagy-lysosome pathway in neurodegeneration associated with Parkinson’s disease. Brain, 2008. 131(Pt 8), 1969-1978.

[105] Luchansky, S.J.; Lansbury, P.T., Jr.; Stein, R.L. Substrate recognition and catalysis by UCH-L1. Biochemistry, 2006. 45(49), 14717-14125.

